# Systematic analysis of *atx*-2 suppressors reveals a novel regulator of PAR-5/14-3-3sigma function during mitosis in *Caenorhabditis elegans*

**DOI:** 10.1101/173856

**Authors:** Megan M. Gnazzo, Alex R. Villarreal, Ahna R. Skop

## Abstract

RNA regulation plays a critical role in mitosis, yet the mechanisms remain unclear. Our lab recently identified that the conserved RNA-Binding Protein (RBP), ATX-2, regulates cytokinesis by regulating the targeting of ZEN-4 to the spindle midzone through a conserved translation regulator, PAR-5/14-3-3sigma (Gnazzo et al., 2016). While co-depletion of ATX-2 and PAR-5 restored ZEN-4 targeting to the spindle midzone, it did not rescue cell division. To identify factors that may work in concert with ATX-2 to regulate cell division, we conducted a two-part, candidate RNAi suppressor and visual screen to identify factors that are important for cell division and also mediate the targeting of ATX-2 to the centrosomes and the spindle midzone. Using this approach, we identified ten genes that suppress the embryonic lethality defect observed in *atx-2* mutant embryos. These ten genes, including *act-2*, *cgh-1*, *cki-1*, *hum-6*, *par-2*, *rnp-4*, *vab-3*, *vhl-1*, *vps-24*, and *wve-*1, all have some role regulating RNA or the cell cycle. Five of these genes (*cgh-1*, *cki-1*, *vab-3*, *vhl-1*, *vps-*24) fail to target ATX-2 to the centrosomes and midzone when depleted. The strongest suppressor of the *atx-2* phenotype is the DEAD-box RNA helicase CGH-1/DDX6, which has been implicated in cell division, RNA processing and translation, and neuronal function. Loss of CGH-1 rescued the cytokinesis defect and also restored ZEN-4 localization to the spindle midzone. ATX-2 and CGH-1 are mutually required for their localization to centrosomes and the spindle midzone. Our findings provide the first functional evidence that CGH-1/DDX6 regulates ATX-2 function during mitosis to target ZEN-4 to the spindle midzone via PAR-5/14-3-3sigma. We suggest that RNA machinery is necessary for the completion of cytokinesis.

## Introduction

The last step in cell division, cytokinesis, is a critical step for daughter cell separation (D'Avino et al., 2015; Green et al., 2012). Daughter cell separation requires coordination of the mitotic spindle, actin cytoskeleton, plasma membrane and cytoplasmic components (D'Avino et al., 2015; Dionne et al., 2015; Mierzwa and Gerlich, 2014). Factors necessary for cell separation localize to the spindle midzone and midbody to mediate cleavage furrow formation and serve as platforms for the spatial and temporal control mechanisms that mediate abscission or cell separation (Glotzer, 2009; Gould, 2016; Pohl, 2017). Abscission and cell proliferation defects often result in multinucleate cells (Jordan et al., 2011; Normand and King, 2010; Sagona and Stenmark, 2010), and have been associated with several human diseases, including Hodgkin lymphoma (Salipante et al., 2009), MARCH syndrome (Frosk et al., 2017) and microcephaly (Basit et al., 2016; Pulvers et al., 2010). Thus, strict regulation of the last steps of cytokinesis is necessary for proper cell division, cell proliferation, and human health.

Recent studies in several systems have discovered that RNA binding proteins, mRNAs, and lncRNAs function and/or localize to the centrosomes, metaphase plate, kinetochores or midbody during mitosis and are suggested to play regulatory roles during the cell cycle (Aviner et al., 2017; Blower et al., 2007; Gnazzo et al., 2016; Skop et al., 2004; Stubenvoll et al., 2016; Yokoshi et al., 2014; Zheng et al., 2010). In *C. elegans*, the RNA-binding proteins CAR-1, SPN-4, LARP-1, and ATX-2 are all required during mitosis (Audhya et al., 2005; Burrows et al., 2010; Gnazzo et al., 2016; Gomes et al., 2001; Squirrell et al., 2006; Stubenvoll et al., 2016), and both CAR-1 and ATX-2 are necessary for spindle midzone assembly, midbody function, and the completion of cell division (Audhya et al., 2005; Gnazzo et al., 2016; Squirrell et al., 2006). In yeast, the DEAD-box RNA helicase, Ded1/DDX3, and the 5’ mRNA cap-binding protein, Cdc33/eI4FE, both mediate translation and cell cycle checkpoints (Brenner et al., 1988; Grallert et al., 2000; Kronja and Orr-Weaver, 2011). It has been recently discovered that Cyclin A2, has a CDK-independent role as an RNA binding protein that modulates the translation of Mre11 mRNA necessary to repair double stranded breaks, suggesting a broader role for cyclins in mediating mitotic events than previously thought (Kanakkanthara et al., 2016). In *Drosophila and Xenopus*, mRNAs and their encoded proteins associate with the mitotic spindle (Blower et al., 2007; Lecuyer et al., 2007). Long non-coding RNAs (lncRNAs) have been found at the spindle midzone and to the midbody (Zheng et al., 2010), and a specific non-coding RNA, LINC00152, is necessary for mitotic progression (Notzold et al., 2017). These data suggest that mRNAs, their encoded proteins, and non-coding RNAs are all necessary factors required during mitosis, but many questions remain, including how these factors are spatially and temporally regulated at each step in the cell cycle. To try to uncover mechanisms, recent mitotic ribosomal profiling and proteomic analysis of polyribosomes revealed about 200 mRNAs that are translated upon mitotic entry (Tanenbaum et al., 2015). Furthermore, the translation machinery itself appears to be regulated in part by hnRNP C during mitosis, such that daughter cells can efficiently synthesize proteins as they enter G1 (Aviner, 2017), the stage at which the midbody is being formed. One factor that appears to mediate translation during cytokinesis is an isoform of 14-3-3, sigma, which binds directly to mitosis-specific factors involved in cap-independent translation. Loss of 14-3-3sigma leads to defects in proper targeting of Polo-like kinase 1 to midbodies and cytokinesis (Wilker et al., 2007). Our lab has shown that the 14-3-3sigma homolog, PAR-5 appears to be regulated during mitosis via a conserved RBP, Ataxin-2/ATX-2 (Gnazzo, 2016). Taken together, proper regulation of RBPs, mRNAs, non-coding RNAs, and translation regulators during cytokinesis is critical for successful cell division, but the mechanisms remain unclear.

To determine how ATX-2 is specifically regulated during mitosis, we utilized a candidate suppression screen to identify factors that suppress embryonic phenotypes observed in *atx-2(ne4297)* temperature sensitive embryos (Gnazzo, 2016). A secondary visual screen was performed to identify candidates that also led to the mis-targeting of the ATX-2 during mitosis. The strongest suppressor of *atx-2(ne4297)* was found to be the conserved DEAD-box RNA helicase CGH-1. CGH-1 belongs to the DDX6 family of helicases (Navarro et al., 2001). DDX6 plays important roles in RNA storage, and translational repression and decay during development (Bourgeois et al., 2016; Ostareck et al., 2014). Moreover, DDX6 interacts with Ataxin-2 in RNA granules to regulate global mRNA translation, stability and degradation (Nonhoff et al., 2007). We determined that ATX-2 and CGH-1 control cell division by coordinately regulating the targeting of a conserved translational regulator, PAR-5/14-3-3sigma and subsequently the key cell division kinesin, ZEN-4/MKLP1/CHO1 to mitotic structures. These data suggest that multiple RBPs work in concert to mediate the spatial and temporal regulation of cell division factors during mitosis.

## Results

### Candidate RNAi suppressor screen identified 10 atx-2 suppressors

To identify mitotic interactors of ATX-2, we conducted a candidate RNAi suppressor screen, an approach that has been very fruitful in uncovering genes involved in various developmental processes in *C. elegans* (Dorfman et al., 2009; Fievet et al., 2013; O'Rourke et al., 2007; Wang and Sherwood, 2011). In this screen, we utilized a temperature-sensitive (ts) allele of *atx-2(ne4297)* that lays 100% dead embryos (0% embryonic viability) at the restrictive temperature (25°C) (Gnazzo et al., 2016). We chose 92 candidate genes (Table 1) that were predicted to interact with ATX-2 based on several criteria: protein localization pattern, involvement in a similar biological process, and similar gene ontology using several bioinformatic databases (Chatr-Aryamontri et al., 2017; Licata and Orchard, 2016; Szklarczyk et al., 2015; Zhong and Sternberg, 2006)(see Methods). To assay *atx-2* suppression, *atx-2(ne4297)* hermaphrodites were grown on candidate RNAi feeding bacteria for 24 hours at 25°C and then scored for embryonic viability (Fig. 1A). Any depletion that reproducibly yielded viable progeny was identified as an *atx-2* suppressor. Ten candidate genes were identified using this strategy (Fig. 1B, Table 2). The strongest *atx-2* suppressor was the human DDX6 ortholog CGH-1, which when knocked down in concert with ATX-2 suppressed the atx-2*(ne4297)* embryonic lethality phenotype (34.53 ± 6.8% embryos survived compared to 0% for atx-2 alone). The next strongest *atx-2* suppressor was VHL-1, the ortholog of the Von Hippel-Lindau tumor suppressor gene VHL, which led to 19.05 ± 11.76% embryonic viability (Fig 1B; Table 2).

**Figure 1.**
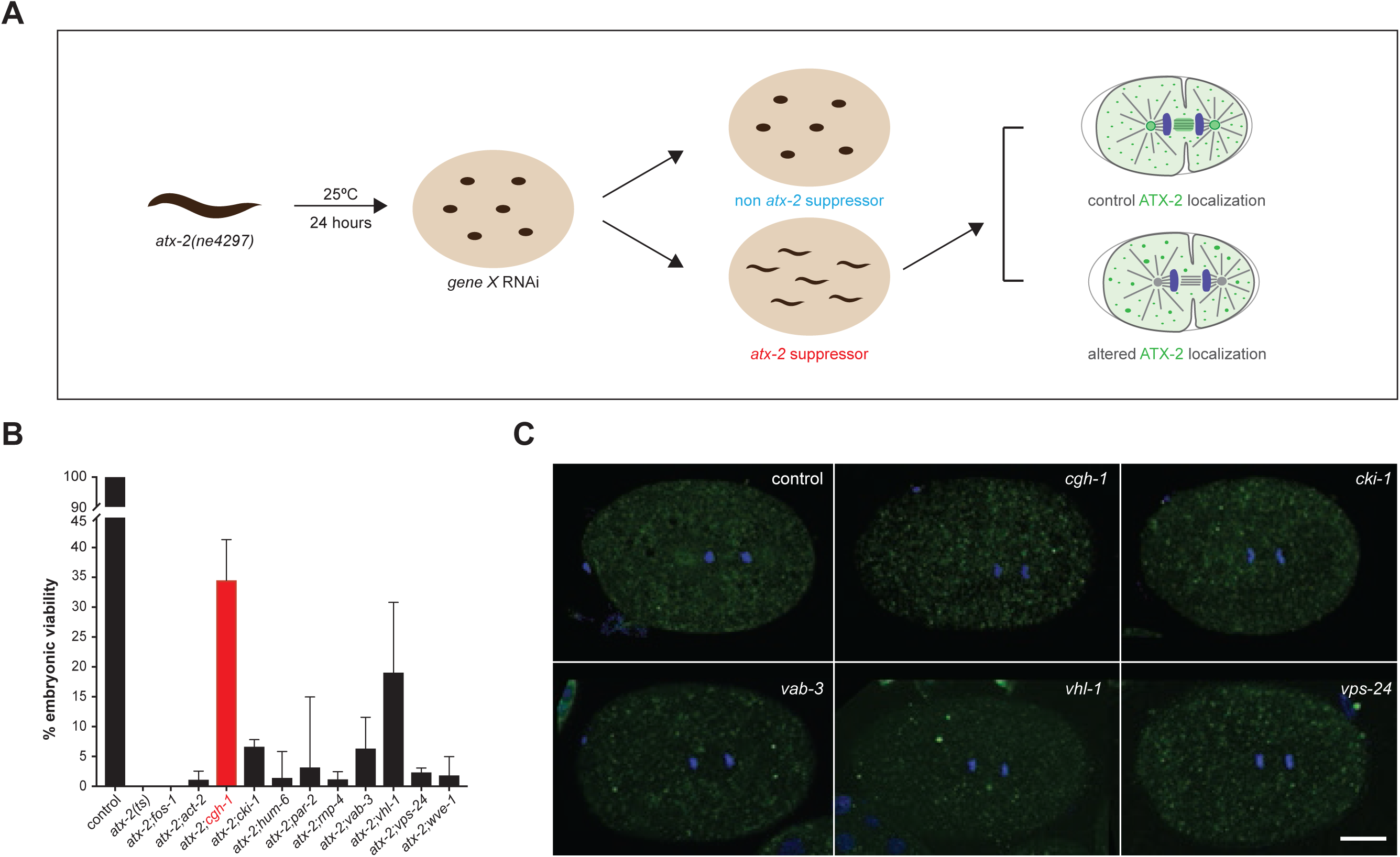
Candidate RNAi suppression screen identifies 10 *atx-2* suppressor genes. A) Schematic showing the candidate RNAi suppression screen conducted. L4 staged *atx-2(ne4297)* worms were placed onto feeding RNAi plates specific to one of 92 candidate genes. Plates were shifted to 25°C while worms laid embryos for 24 hours before removing the adult. After 24 hours, plates were scored for the presence of hatched embryos. Any candidate gene with hatched embryos was considered an *atx-2* suppressor, while any candidate gene with 100% dead embryos was not considered to be an *atx-2* suppressor. *atx-2(ne4297)* worms lay 100% dead embryos at 25°C. B) Ten candidate *atx-2* suppressor genes were identified. CGH-1 was the strongest suppressor with 34.53 ± 6.8% embryonic viability, and is highlighted in red. The second strongest suppressor was VHL-1 with 19.05 ± 11.76% embryonic viability. Control/N2, *atx-2(ne4297) and fos-1* were included as experimental controls. C) The mitotic localization of ATX-2 to the centrosomes, spindle midzone and cytoplasmic puncta was altered by depletion of candidate genes. Fixed and immunostained ATX-2-GFP embryos depleted with fRNAi for *cgh-*1, cki*-1*, *vab-3, vhl-1*, and *vps-24 are shown*. All candidate suppressors alter the localization of ATX-2 to the centrosome, midzone and cytoplasmic puncta.

**Table 1.**
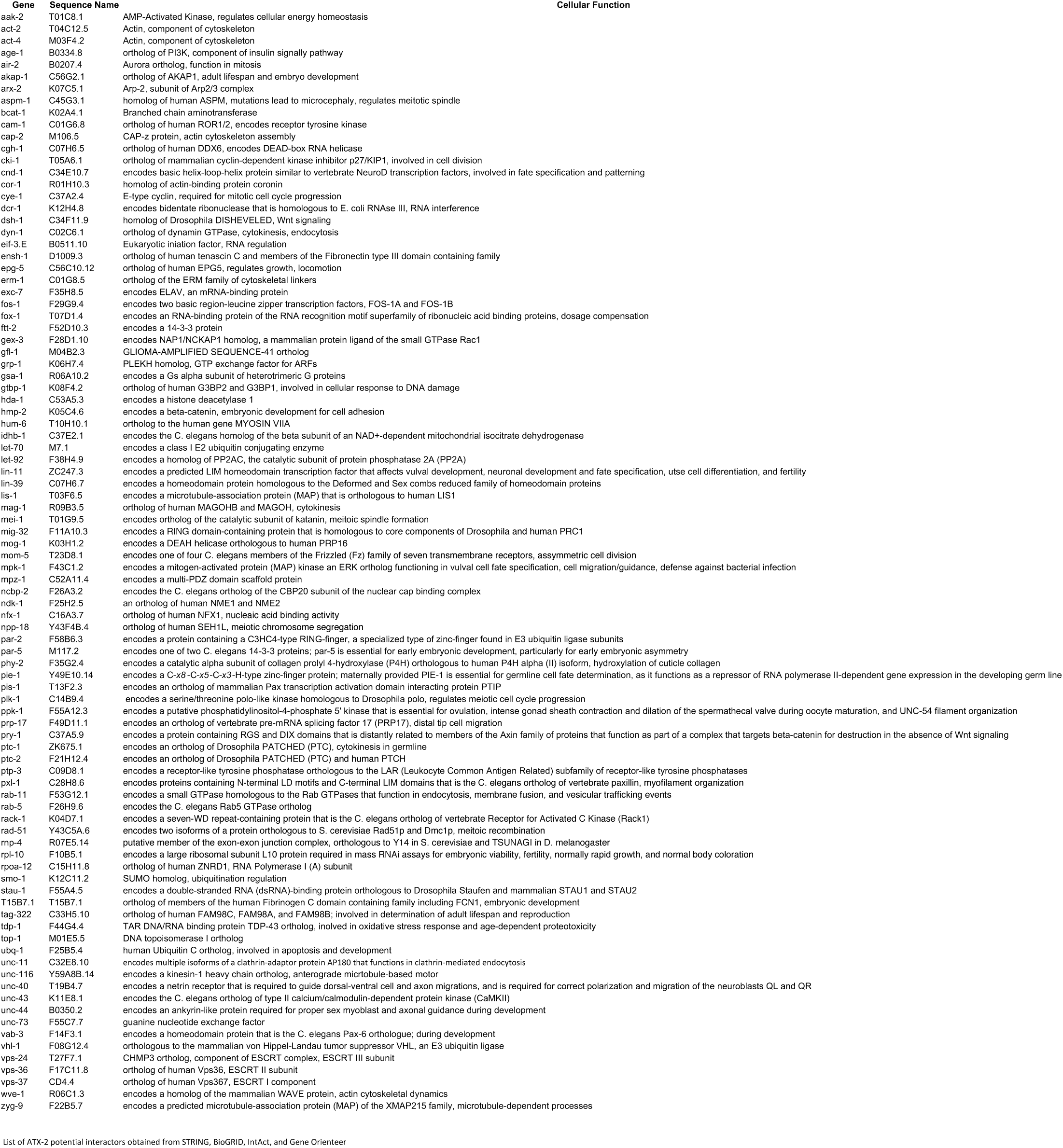
The 92 candidate genes screened in this study. Table of the 92 candidate genes selected to screen in this study. Genes were chosen based on predicted interaction with ATX-2 using STRING, BioGRID, IntAct, and Gene Orienteer (Chatr-Aryamontri et al., 2017; Licata and Orchard, 2016; Szklarczyk et al., 2015; Zhong and Sternberg, 2006). Genes are listed by *C. elegans* gene and sequence name with associated cellular function obtained from Wormbase.

**Table 2.**
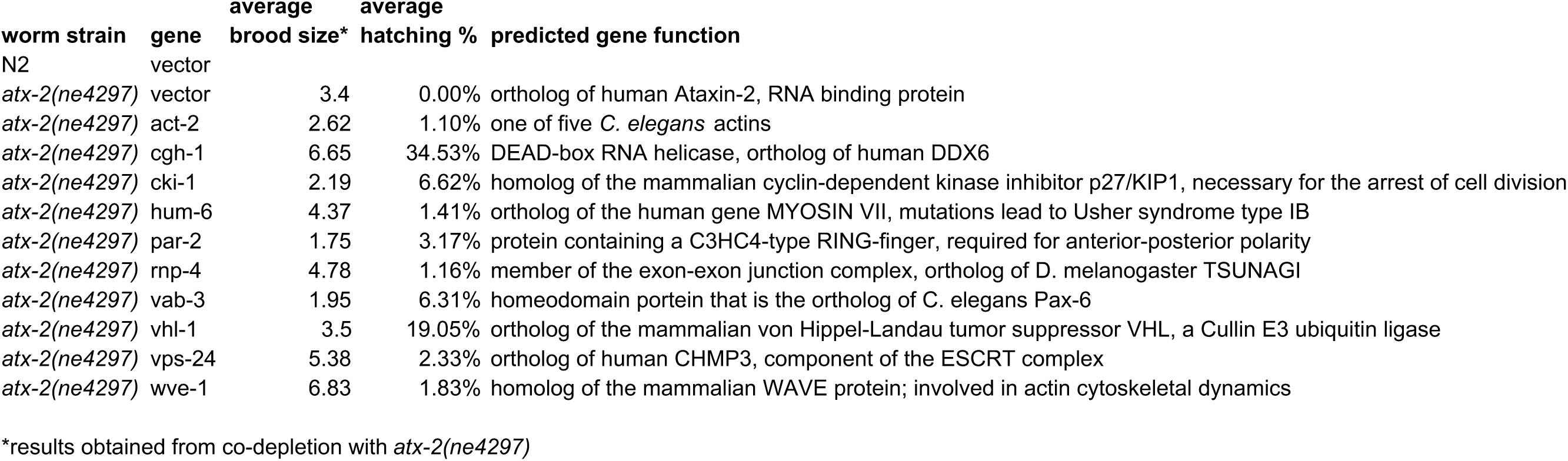
The 10 identified suppressors in the *atx-2(ne4297)* RNAi screen. Table of the 10 identified *atx-2* suppressors. Each gene reproducibly yielded >0% embryonic viability when co-depleted with ATX-2 at 25°C. Average brood size was calculated as number of embryos laid within first 24-hour period of placing *atx-2(ne4297)* on RNAi plates. Average hatching percentage was calculated as number of hatched embryos divided by total number of embryos laid multiplied by 100. Predicted gene function was obtained from the WormBase and UniProt (The UniProt, 2017).

To focus our analysis on genes with specific roles in mitosis, we employed a secondary visual screen to determine if the strongest candidate suppressors altered the mitotic localization of ATX-2 (Gnazzo et al., 2016). To do this, we fixed and immunostained ATX-2-GFP-expressing embryos that were treated with vector control, *cgh-1*, *cki-1*, *vab-3*, *vhl-1*, or *vps-24* fRNAi. These five candidates were chosen because they were consistently the strongest *atx-2(ne4297)* suppressors and yielded the highest percentage embryonic viability when co-depleted with ATX-2. In control embryos, ATX-2-GFP localized to distinct puncta throughout the cytoplasm and accumulated at the centrosomes and spindle midzone during anaphase ((Gnazzo et al., 2016); Fig. 1C). Single depletions of CGH-1, CKI-1, VAB-3, VHL-1, or VPS-24 all disrupted the spindle and midzone localization of ATX-2-GFP, suggesting that each suppressor regulates the mitotic localization of ATX-2 (Fig. 1C). In each case, ATX-2-GFP localization to cytoplasmic puncta was still observed; however, the patterning and size of the ATX-2-GFP puncta was altered in each condition. For example, in *cgh-1* fRNAi, *vab-3* fRNAi-, and *vhl-1* fRNAi-treated embryos, the puncta were larger and less numerous in the cytoplasm (Fig. 1C). Overall, we identified five *atx-2(ne4297)* suppressors that suppressed the lethality of *atx-2(ne4297)* and also altered the size and distribution of ATX-2-GFP-containing puncta during mitosis. Given the screen results, the highest percent embryonic viability and large, aggregated ATX-2-GFP puncta observed in CGH-1 depleted embryos, further analysis focused on understanding the interaction between CGH-1 and ATX-2.

#### CGH-1, a conserved DEAD-box helicase, suppresses the ATX-2 cell division defect

To begin our analysis, we utilized Differential Interference Contrast (DIC) time-lapse microscopy to visualize the first cell division in early *C. elegans* embryos in control and *cgh-1* fRNAi-treated embryos. In control embryos, the cleavage furrow completed to form two blastomeres of unequal size (Fig. 2C: control; Movie S1). In single depletion *cgh-1* fRNAi-treated embryos, cell division failed in 28.57% (n=2/7) of embryos. Here, the cleavage furrow initiated (8:30min), completed (11:10min) and subsequently retracted (22:20min), resulting in a multinucleate embryo (Fig. 2C; S2). This failure rate is similar to that observed in ATX-2 depleted embryos, where cell division failed in 37.50% (n=3/8) of embryos (Fig. 2C; S3) (Gnazzo et al., 2016). Co-depletion of both CGH-1 and ATX-2 suppresses the cell division defects observed in *atx-2(ne4297)* embryos, in 36.36% (n=4/11) of the *atx-2(ne4297)* embryos (Fig. 2C; Movies S4-S5), suggesting that CGH-1 may play an important role in regulating cell division with ATX-2. These findings were not an artifact of the full length *cgh-1* fRNAi used, as two additional RNAi feeding constructs made to target either the N or C terminus of CGH-1 also rescued the *atx-2(ne4297)* defects (Fig. S1).

**Figure 2.**
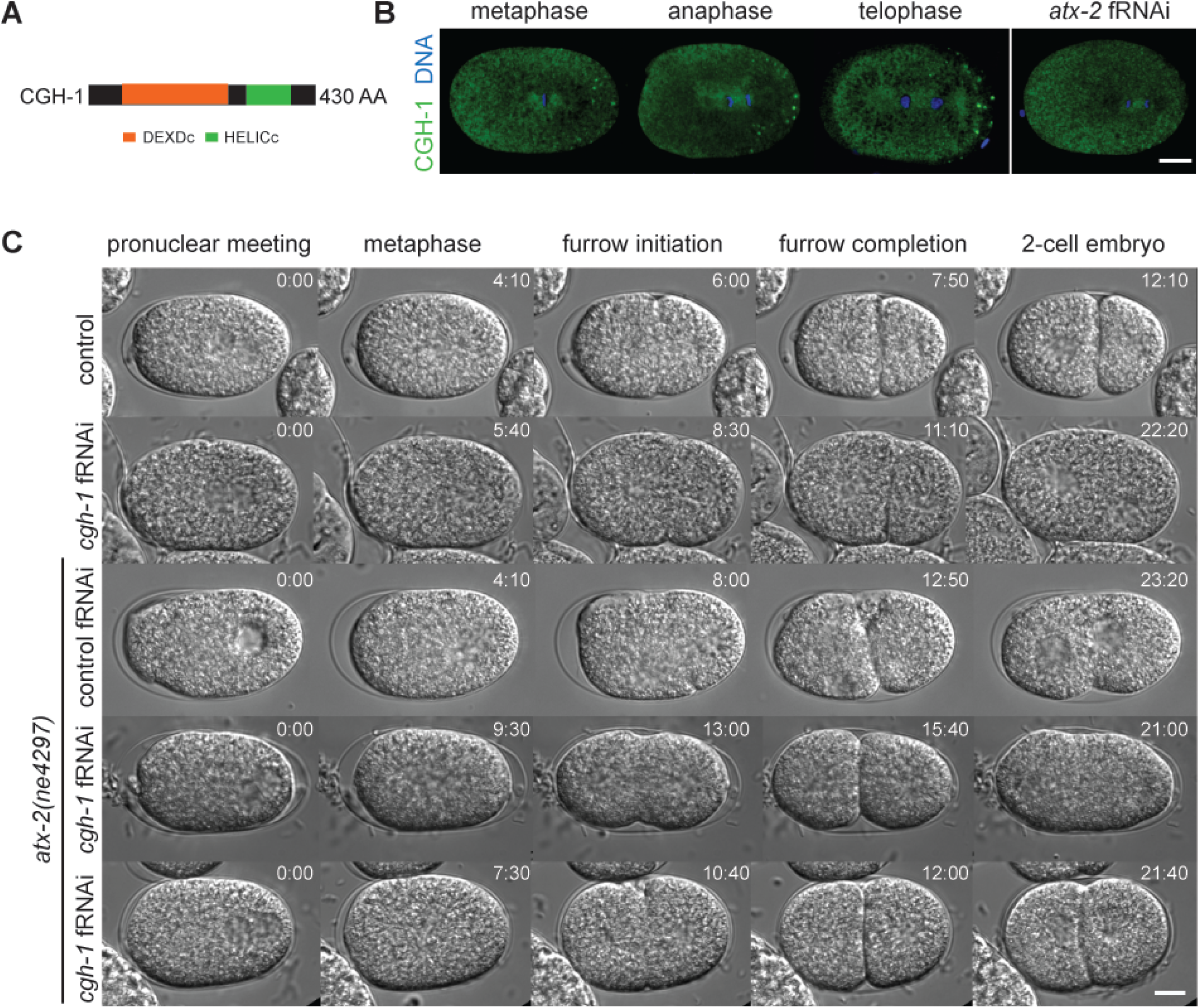
*cgh-1* RNAi suppresses *atx-2(ne4297)*. A) The CGH-1 protein has two conserved domains: DEXDc and HELICc. DEXDc is a DEAD-like helicase domain that is involved in ATP-dependent RNA binding, and HELICc is a helicase c-terminal domain that is associated with DEXDc. In CGH-1 both domains function to unwind RNA. B) CGH-1 localization in control and ATX-2 depleted embryos. In control embryos stained and fixed with anti-CGH-1, CGH-1 localized around the metaphase plate, the spindle midzone and to P-granules. In ATX-2 depleted embryos, the centrosome localization was lost and the P granule localization was reduced. However, the spindle midzone localization was maintained. Scale bar, 10 μm. C) CGH-1 depletion rescues the cytokinesis defects observed in ATX-2 depleted embryos. Differential Interference Contrast (DIC) time-lapse images of control, *cgh-1* fRNAi-treated, *atx-2(ne4297)*, and *atx-2(ne4297)*; *cgh-1* fRNAi-treated embryos throughout the first cell division. In control embryos, the cleavage furrow initiates (6:00) and subsequently completes (7:50), resulting in a 2-cell embryo (12:10). In *cgh-1* fRNAi-treated embryos, the cleavage furrow initiates (8:30), completes (11:10) and subsequently retracts (22:20), resulting in cytokinesis failure and a multinucleate embryo. In *atx-2(ne4297)* embryos, the cleavage furrow initiates (8:00) and subsequently retracts, resulting in a multinucleate embryo (24:00). When *atx-2(ne4297)* embryos are treated with *cgh-*1 fRNAi, cytokinesis is rescued in 36.36% of the time. Here, the furrow initiates (10:40) and subsequently completes (12:00), resulting in a 2-cell embryo (21:40). In instances were CGH-1 depletion does not suppress *atx-2(ne4297)*, the cleavage furrow initiates (13:00) and subsequently retracts, resulting in a multinucleate embryo (24:20). Times, in min:sec, are given relative to pronuclear meeting. Scale bar, 10 μm.

Next, we sought to determine if the CGH-1 shared a similar cell cycle localization pattern to ATX-2. Previously, CGH-1 has been shown to localize to P granules in the posterior of the early embryo (Audhya et al., 2005; Boag et al., 2005; Navarro et al., 2001); however, the temporal and spatial localization pattern during the cell cycle had not been explored. Here, control and *atx-2* fRNAi-treated embryos were fixed and stained with anti-CGH-1 antibody and visualized using confocal microscopy. During the cell cycle, CGH-1 was generally cytoplasmic with and enrichment around the mitotic spindle and P granules. Here, CGH-1 localized in a cloud around the metaphase plate and was found at the centrosomes on the spindle midzone in anaphase (Fig. 2B). In *atx-2* fRNAi-treated embryos, the centrosomes and P granule localization were lost, but the spindle midzone localization remained, suggesting that ATX-2 is only necessary to maintain CGH-1 at centrosomes and P granules. Given that the spindle midzone localization of CGH-1 was unaffected by ATX-2 depletion, CGH-1 targeting to the midzone likely occurs independently of ATX-2.

### CGH-1 depletion rescued cell division defects observed in *atx-2(4297)* embryos

Given that ATX-2 and CGH-1 function to regulate cell division, we wanted to determine if CGH-1 regulated the necessary cell division pathway components, ZEN-4 and PAR-5 (Basant et al., 2015; Douglas et al., 2010; Gnazzo et al., 2016). ZEN-4 is a component of the Centralspindlin complex and its localization to the spindle midzone is essential for cell division to complete (Raich et al., 1998). PAR-5 is a 14-3-3sigma ortholog prevents ZEN-4 from localizing to the spindle midzone (Basant et al., 2015; Douglas et al., 2010). We previously demonstrated that ATX-2 negatively regulates the protein expression of PAR-5 and the proper targeting of ZEN-4 to the spindle midzone (Gnazzo et al., 2016). To determine if CGH-1 suppressed the observed *atx-2* embryonic defects, we utilized live cell imaging and the PAR-5-GFP (Mikl and Cowan, 2014), and ZEN-4-GFP (Kaitna et al., 2000) worm strains. In control embryos, PAR-5-GFP localized in a cell cycle dependent manner around the chromatin during Nuclear Envelope Break Down (NEBD), in a cloud around metaphase plate, centrosomes throughout the cell cycle (black arrowheads), the spindle midzone in anaphase, faintly along the cleavage furrow and at the midbody in telophase (white arrow head)(Fig. 3A; Movie S6; (Gnazzo et al., 2016)). In *atx-2* fRNAi-treated embryos, PAR-5-GFP was enriched around chromatin during NEBD, at the centrosomes from metaphase to telophase (black arrowheads), and specifically during anaphase, PAR-5-GFP puncta dispersed from the spindle midzone into the cytoplasm In telophase, chromatin associated PAR-5-GFP is lost and PAR-5-GFP accumulated at the periphery of the centrosomes (Fig. 3A; Movie S7; (Gnazzo et al., 2016)). CGH-1 depletion did not alter PAR-5-GFP localization to the metaphase plate, centrosomes (black arrowheads) or cleavage furrow. However, in *cgh-1* fRNAi-treated embryos PAR-5-GFP was lost from the spindle midzone and had fewer PAR-5-GFP labeled P granules (Fig. 3A; Movie S8). CGH-1 may specifically regulate PAR-5 localization at the spindle midzone and P granules, consistent with the previously defined functions of CGH-1 (Audhya et al., 2005; Boag et al., 2005; Updike and Strome, 2009). Lastly, co-depletion of ATX-2 and CGH-1 rescued the PAR-5-GFP defects observed in ATX-2-depleted embryos, restoring PAR-5-GFP localization to the spindle midzone, the periphery of the centrosomes (black arrowheads), and P granules (Fig. 3A).

**Figure 3.**
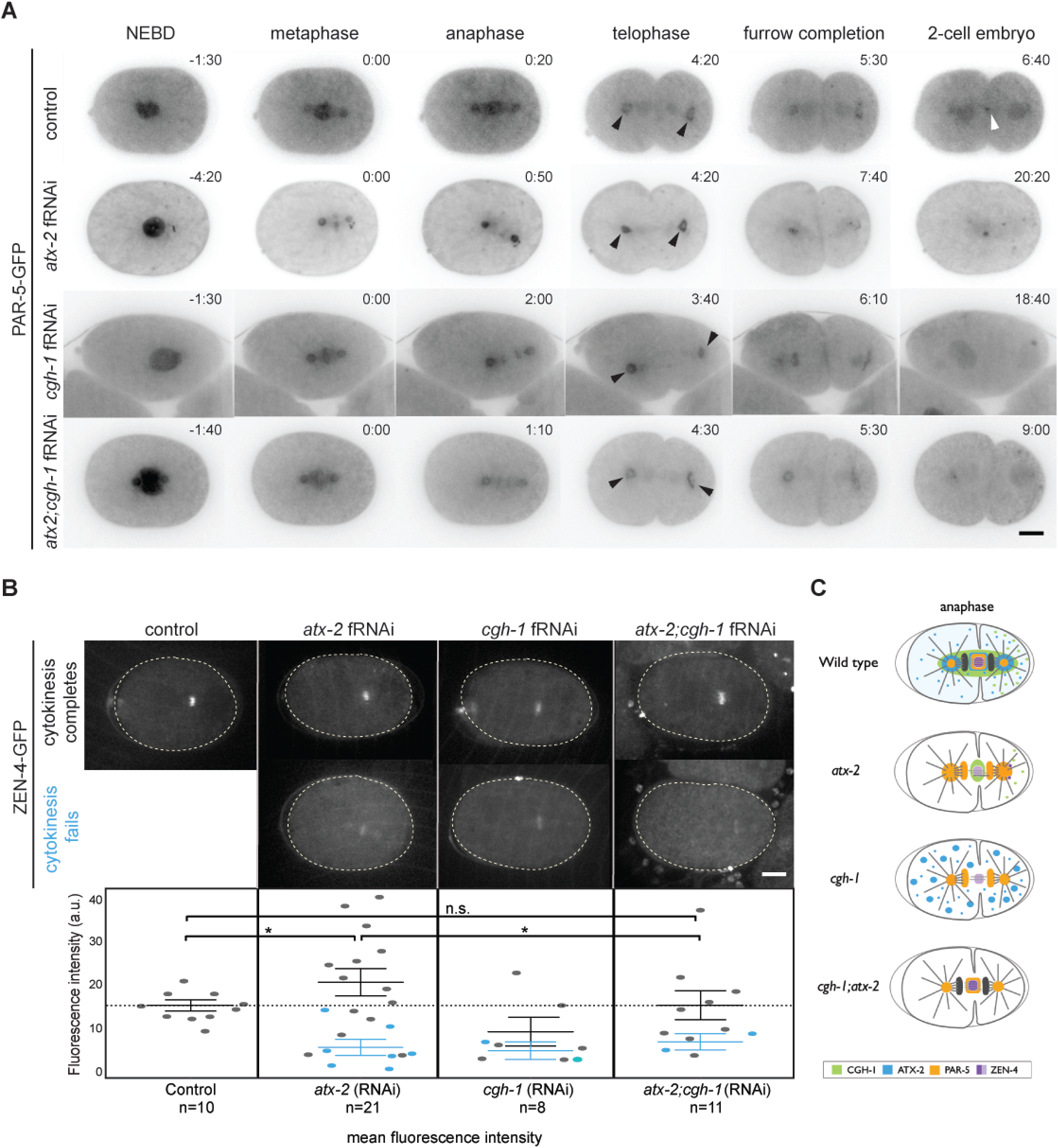
PAR-5-GFP and ZEN-4-GFP dynamics in *atx-2;cgh-1* embryos. A) PAR-5-GFP dynamics in control, *atx-2* fRNAi-, *cgh-1* fRNAi-, and *atx-2;cgh-1* fRNAi-treated embryos. In control embryos, PAR-5-GFP localizes to nuclei (-1:30min), the metaphase plate (0:00min), centrosomes and the spindle midzone (4:20min, black arrowheads), faintly along the cleavage furrow and posterior cytoplasmic granules (5:30), and the midbody (6:40min, white arrowhead). In *atx-2* fRNAi-treated embryos, PAR-5-GFP accumulates at the metaphase plate (0:00min) and at the centrosomes (0:50min, black arrowheads). PAR-5-GFP then associates with the spindle midzone and midbody, and faintly at the cleavage furrow (7:40min). In *cgh-1* fRNAi-treated embryos, PAR-5-GFP localizes to nuclei (-1:30 min), the metaphase plate (0:00 min), centrosomes (3:40 min, black arrowheads), and the cleavage furrow (6:10 min). Co-depletion of ATX-2 and CGH-1 suppresses the PAR-5-GFP localization defects observed in ATX-2 depleted embryos. PAR-5-GFP localizes to the metaphase plate (0:00 min), the spindle midzone and centrosomes (4:30 min, black arrowheads), the cleavage furrow (5:30 min), and posterior cytoplasmic granules (9:00 min). Times, in min:sec, are given relative to metaphase. Scale bar, 10 μm. B) Visualization of ZEN-4-GFP localization in control, *atx-2* fRNAi-, *cgh-1* fRNAi-, and *atx-2; cgh-*1 fRNAi-treated embryos. In control embryos, ZEN-4 localizes to the spindle midzone. In *atx-2* fRNAi- and *cgh-1* fRNAi-treated embryos ZEN-4 expression at the midzone is reduced. In *atx-2; cgh-*1 fRNAi-treated embryos, ZEN-4-GFP expression at the midzone is rescues. Images are maximum projections of 0.5 μm Z-stacks spanning 1.5 μm. Scale bars, 10 μm. Quantification of ZEN-4-GFP fluorescence intensity in control, *atx-2* fRNAi-, *cgh-1* fRNAi-, *and atx-2; cgh-*1 fRNAi-treated embryos. Each point represents a fluorescence intensity measurement from a single embryo. Grey points represent embryos in which cytokinesis successfully completed; blue points represent embryos in which cytokinesis failed to complete. When cytokinesis fails in *atx-2* fRNAi-treated embryos, ZEN-4-GFP expression is reduced at the spindle midzone. Co-depletion of ATX-2 and CGH-1 rescues the ATX-2 single depletion phenotype. Solid lines denote the mean. Results are the mean ± SEM. C) Summary of ATX-2 and CGH-1 function in cell division regulation. In control cells, ATX-2 and CGH-1 function together to regulate the proper targeting of PAR-5 and CGH-1 to mitotic structures, allowing for successful cell division. In *atx-2* mutants, PAR-5 expression is increased at mitotic structures, ZEN-4 expression is reduced at the spindle midzone, and CGH-1 expression is lost at the mitotic spindle. In *cgh-1* mutants, PAR-5 expression at the spindle is perturbed, ZEN-4 reduced at midzone, and ATX-2 forms into larger and less dense aggregates in the cytoplasm. Loss of both ATX-2 and CGH-1 rescues proper PAR-5 and ZEN-4 expression at mitotic structures and cell division is restored.

Next, we sought to determine if co-depletion of ATX-2 and CGH-1 rescued ZEN-4 localization to the spindle midzone. In control embryos, ZEN-4-GFP localized to the spindle midzone during anaphase and telophase (Fig. 3B), and the fluorescence intensity during anaphase was on average 14.86 a.u. (arbitrary units) (n=10) (measured by drawing a region of interest (ROI) around the spindle midzone in late anaphase). As we previously showed, ATX-2 depletion resulted in improper targeting of ZEN-4-GFP, resulting in decreased expression at the spindle midzone (Fig. 3B; (Gnazzo et al., 2016)). Here, we confirmed that loss of ATX-2 resulted in a significant reduction of ZEN-4-GFP at the midzone compared to control (5.66 a.u. when cytokinesis failed; p value<0.05). Since CGH-1 depletion resulted in cell division failure, we predicted that ZEN-4-GFP localization to the spindle midzone would also be reduced. We found that when cell division failed, less ZEN-4-GFP appeared to target the spindle midzone, yet when quantified, this was not significant (Fig. 3B). To determine if co-depletion of ATX-2 and CGH-1 not only rescues cytokinesis but restores ZEN-4-GFP targeting to the spindle midzone, we measured ZEN-4-GFP fluorescence intensity in the *atx-2;cgh-1* fRNAi-treated embryos (13.93 a.u.; Fig. 3B). This result was significantly different than depleting ATX-2 alone (p value<0.05), suggesting that co-depletion of ATX-2 and CGH-1 rescued ZEN-4-GFP localization to the spindle midzone when cell division was also rescued. Overall, loss of both of CGH-1 and ATX-2 rescued the cell division defects, and the localization of PAR-5-GFP and ZEN-4 to mitotic structures (Fig. 3C). These data suggest that the mitotic RBPs, ATX-2 and CGH-1, play necessary roles in mediating the temporal and spatial localization of critical cell division factors, ZEN-4 and PAR-5. Ultimately ensuring that cell division and daughter cell separation is successful.

### Discussion

The work presented here identified novel genetic factors that function during cell division with the conserved RNA binding protein, ATX-2. We identified ten genes that suppress the embryonic lethality defects observed in *atx-2(ne4297)* embryos. Each of these genes, when co-depleted using feeding RNAi in *atx-2(ne4297)* worms at 25°C, reproducibly yielded greater than 0% embryonic viability, suggesting that they may function in conjunction with ATX-2 to regulate cell division. Of the strongest suppressors, CGH-1 depletion rescued the cell division defects observed in *atx-2(ne4297)* embryos. Further analysis determined that PAR-5 and ZEN-4 targeting to the mitotic spindle or spindle midzone was also restored after co-depleting both CGH-1 and ATX-2. Together, our study suggests that ATX-2 and CGH-1 regulate cell division by mediating the targeting of PAR-5 and ZEN-4 to the spindle.

### Ten genes suppress the *atx-2(ne4297)* cell division defect phenotype

Prior to this study, ATX-2 was found to directly bind to the Poly-A Binding protein, PAB-1, and interact with the germline regulators, TRA-2, GLD-1, GLP-1, and MEX-3 (Ciosk et al., 2004; Maine et al., 2004). However, these interactions do not readily explain why ATX-2 is necessary for normal cell division. The goal of this study was to identify factors that suppressed the cell division phenotype observed in *atx-2(ne4297)* embryos, and thus gain mechanistic insight into the function of ATX-2 in mitosis. Our screen identified ten genes that suppress the cell division defect observed in ATX-2 depleted embryos, including *act-2*, *cgh-1*, *cki-1*, *hum-6*, *par-2*, *rnp-4*, *vab-3*, *vhl-1*, *vps-24*, and *wve-1*. The genes, *act-2*, *cki-1*, *hum-6*, *par-2*, *vps-24*, and *wve-1*, encode proteins that are known to function in the regulation of the cell cycle, cell division or the cytoskeleton (Baker and Titus, 1997; Files et al., 1983; Hong et al., 1998; Kemphues et al., 1988; Kim et al., 2011; Sawa et al., 2003). The remaining genes (*cgh-1*, *rnp-4*, *vab-3*, and *vhl-1)* encode proteins that regulate development or RNA expression, suggesting a larger role for RNA regulation in cell division than was previously appreciated (Bishop et al., 2004; Chamberlin and Sternberg, 1995; Kawano et al., 2004; Navarro et al., 2001). More in depth exploration into the interaction of ATX-2 and these genes will shed light on the role of ATX-2 and how RNA is regulated during cell division and provide insight into human disease.

### CGH-1 and ATX-2 overlap in localization and function

CGH-1 was the strongest suppressor identified in our analysis. In *C. elegans*, CGH-1 is a DEAD-box RNA helicase that is important for cell division animal fertility (Audhya et al., 2005; Boag et al., 2005; Navarro et al., 2001). The human orthologs of CGH-1 and ATX-2 function together in RNA granules to regulate mRNA translation, stability, and degradation (Nonhoff et al., 2007). Additionally, CGH-1 and ATX-2 have a common interaction partner in PAB-1, the Poly-A Binding Protein homolog (Ko et al., 2013), and similar cell division defects in the early embryo (Audhya et al., 2005; Gnazzo et al., 2016), suggesting a broad role in mRNA metabolism during mitosis. CGH-1 directly binds to PAB-1 in P granules, and both CGH-1 and ATX-2 are modifiers of P granule localization and dynamics (Gnazzo et al., 2016; Hubstenberger et al., 2015; Updike and Strome, 2009; Wood et al., 2016), indicating that all of these RPBs play critical roles in RNA dynamics during asymmetric cell division events. In this work, we found that ATX-2 and CGH-1 function together to regulate cell division through the proper targeting of PAR-5 and ZEN-4 to the spindle midzone. We identified that both ATX-2 and CGH-1 localize to mitotic structures during cell division, including the around the metaphase plate, centrosomes, spindle midzone, furrow and midbody (Fig. 1C, 2B), suggesting that ATX-2 and CGH-1 may regulate cell division by regulating RNA on mitotic structures.

### Novel interactors of ATX-2 and implications for human disease

The identification of several novel interactors of ATX-2 has implications for understanding the neurodegenerative disease and the formation of neuroblastomas (Ostrowski et al., 2017). Expansion of the polyglutamine (polyQ) domain leads to age-related motor, Purkinje, and basal ganglia neuronal atrophy in SCA2 and ALS patients (Gierga et al., 2005; Rub et al., 2003). Ataxin-2 normally relocates to the stress granules in periods of stress to maintain the health of neurons, but in these patients this function is abrogated (Auburger et al., 2017; Kaehler et al., 2012). In mouse models, Ataxin-2 overexpression increases the severity of the neuronal atrophy, but loss of Ataxin-2 postpones the disease state and extend life span (Becker et al., 2017). Interesting, tumor growth regression coincides with the upregulation of Ataxin-2 in neuroblastoma tumors (Auburger et al., 2017), suggesting that Ataxin-2 may play a general role in modulating cell cycle, as we have shown here. The human ortholog of CGH-1, DDX6, although not a novel interactor, functions in neurons in stress granules with Ataxin-2 as a posttranscriptional regulator (Nonhoff et al., 2007), and also in mediating the temporal and spatial function of miRNAs to maintain the balance between stem cell maintenance and neuronal differentiation (Coolen et al., 2012; Nicklas et al., 2015; Palm et al., 2013). Ataxin-2 and DDX6 could affect global RNA levels particularly along microtubule dense regions especially at growth cones or the synapse (Gage, 2000; Sun et al., 2013), or affect the process of neurogenesis in the adult brain in the hippocampus, which is particular important for normal brain function in mice, songbirds, monkey and humans (Eriksson et al., 1998; Nicklas et al., 2015; Reynolds and Weiss, 1992; Spalding et al., 2013). Failures in cell division during neurogenesis could lead to less neurons, particularly in these parts of the brain, causing muscle paralysis or lead to neuroblastomas given that multinucleated neurons or microglia potentially lead to neurodegeneration as observed in Amyotrophic lateral sclerosis (ALS) or the progression of neuroblastomas (Fendrick et al., 2007; Nicklas et al., 2015). Lastly, several novel interactors for ATX-2 in cell division were identified, including the ESCRT III complex component, VPS-24, and the Von-Hippel-Lindau (VHL) tumor suppressor homolog, VHL-1, both proteins important for cell cycle progression, membrane remodeling, mitosis and cytokinesis (Bhutta et al., 2014; Kaelin, 2007; Kim et al., 2011; Schuh and Audhya, 2014) Recent work has shown that the ESCRT II complex, which can act with ESCRT III but also independently, can control axon membrane growth and local protein synthesis (Filip, 2016), indicating Ataxin-2 could modulate membrane dynamics by balancing the levels of protein synthesis locally in neurons. Loss of human VHL is often associated with renal carcinomas but also central nervous system blastomas (Kaelin, 2002), suggesting that Ataxin-2 could function in concert with VHL in the nervous system to modulate neurogenesis, for example. It will be particularly interesting to determine the mechanisms of DDX6/CGH-1, ESCRT III/VPS-24, and VHL/VHL-1 function in the brain given the connections to cell division and formation of neuroblastomas. Further exploration into the mechanism of interaction between ATX-2 and these novel factors will provide additional insight into the pathology of SCA2, microcephaly, and several types of brain tumors. Overall, the early *C. elegans* embryo provides an excellent model for understanding and identifying novel molecular pathways by which ATX-2/Ataxin-2 functions in the context of human disease and roles where cell division plays a critical role.

## Materials and Methods

### Worm strains

The following worm strains were used: N2 wild type (Brenner, 1974), WM210 (*atx-2(ts)*; ne4297; (Gnazzo et al., 2016)), MAD63 (ATX-2-GFP; (Gnazzo et al., 2016)), MG170 (ZEN-4-GFP; (Kaitna et al., 2000)), UE50 (GFP-PAR-5; (Mikl and Cowan, 2014)).

### *atx-2* suppression screen

The *atx-2* suppression screen was performed using WM210 (*atx-2(ts)*; ne4297) (Gnazzo et al., 2016) worms at the non-permissive temperature (25°C). Feeding RNAi plates were seeded with sequence verified, bacterial cultures (100uL) from the Ahringer feeding library (Kamath et al., 2003). Genes were chosen based on predicted interaction with ATX-2 using STRING, BioGRID, IntAct, and Gene Orienteer (Chatr-Aryamontri et al., 2017; Licata and Orchard, 2016; Szklarczyk et al., 2015; Zhong and Sternberg, 2006). The empty feeding vector, pL4440, was used as a negative control with N2 worms and as a positive control with *atx-*2*(ne4297)* worms. One L4 hermaphrodite was placed on a seeded RNAi plate at 25°C for 24 hours to lay embryos, before subsequent adult worm removal. 24 hours after worm removal, the embryos were counted and scored to determine percentage of embryonic viability. Since, *atx-* 2*(ne4297)* control worms lay 100% dead embryos, any candidate gene with reproducible embryonic viability >0% was considered a suppressor of *atx-2*.

### RNA interference

ATX-2, CGH-1, CKI-1, VAB-3, VHL-1, and VPS-24 depletions were performed using the RNAi pL4440 feeding vectors from the Ahringer feeding RNAi library (Ahringer, 2006; Kamath and Ahringer, 2003; Kamath et al., 2003; Kamath et al., 2001; Timmons and Fire, 1998). Feeding RNAi plates were seeded with 100uL of bacterial culture and allowed to grow overnight. L4 hermaphrodites were placed on seeded RNAi plates for 18-22 hours at 24°C before imaging. CGH-1 N-terminus and C-terminus feeding RNAi vectors were constructed using the previously published protocol (Kamath and Ahringer, 2003). To make the CGH-1 N-terminus (1-166AA) feeding RNAi vector the following primers were used ATGAGTGGAGCGGAGCAAC (fwd) and ATGACCAAGTGAACCGTTCC (rev). GGAATTCTCGACCGCCTC (fwd) and TAAGCAGTGGTCTCATCG (rev) were used to target the C-terminus of CGH-1 (223-430AA).

### Live-imaging

Embryo dissection was performed in Shelton’s Growth Media and embryos were mounted on 2% agar pads (made with Egg salt buffer) in another drop (20uL) of Shelton’s Media (Shelton and Bowerman, 1996). The agar pads were covered with a 22mm x 22 mm coverslip and sealed in place with Vaseline. We have found that this modified mount abrogates the osmotic defect observed in *atx-2* fRNAi or *atx-2(ne4297)* embryos while allowing for optimal imaging during the first two cell divisions. Time-lapse images were taken every 10 seconds using a 200M inverted Axioscope microscope (Carl Zeiss) equipped with a spinning disk confocal scan head (QLC100; VisiTech, Sunderland, United Kingdom), an Orca 285 differential interference contrast (DIC) camera (Hamamatsu), and an Orca ER camera (Hamamatsu). The DIC and Orca ER cameras were operated through MetaMorph software (version 7.7.11.0 Molecular Devices, LLC, Sunnyvale, CA, USA). Image rotating and cropping was performed using Fiji (ImageJ) software (Schindelin et al., 2012).

### Immunostaining

Immunostaining was performed using a methanol-4%formaldehyde fix as described previously (Skop and White, 1998) using the following primary antibodies: anti-GFP rabbit polyclonal (1:100; ab6556; Abcam and anti-CGH-1 rabbit polyclonal (1:100; (Navarro et al., 2001)) diluted in PBSB (1X PBS, 1% BSA) and incubated overnight at 4°C. Unbound primary antibodies were removed by washing three times for 5 minutes each with PBST (1X PBS, 0.5% Tween). Secondary antibodies used were applied as follows: Alexa Fluor 488 anti-mouse (A-11001; Molecular Probes) and Alexa Fluor 488 anti-rabbit (A-11008; Molecular Probes) with all secondary antibodies diluted 1:200 in PBSB. Following 2 hours of incubation in the dark at ambient conditions, unbound secondary antibodies were removed as above. The fixed and stained embryos were mounted with 8μL of Vectashield with DAPI (H-1200; Vector Laboratories). Confocal imaging was performed on a departmental Zeiss 510 Confocal LSM operated with ZEN software (Carl Ziess, Germany).

## Acknowledgements

We thank Peter Boag, Erik Jessen, Michael Glotzer, Andy Golden, Kevin O’Connell and Carrie Cowan for the generous gifts of reagents and guidance. We thank Randall Dahn, Phil Anderson, Kate O’Connor-Giles, Francisco Pelegri, Aki Ikeda, Jen Gilbert for suggestions and advice. We especially thank Wormbase and the CGC. Strains are available upon request. Table 1 contains detailed descriptions of all genes screened in this study with the sequence name available at Wormbase. This work was supported by grants to A.R.S. from the National Science Foundation (MCB 115800), and the *Caenorhabditis* Genetics Center (CGC)(St. Paul, MN), which is funded by the National Institutes of Health Office of Research Infrastructure Programs (P40 OD010440), and WormBase which is supported by grant #U41 HG002223 from the National Human Genome Research Institute at the US National Institutes of Health, the UK Medical Research Council and the UK Biotechnology and Biological Sciences Research Council.

## Supplemental Figure

**Figure S1.**
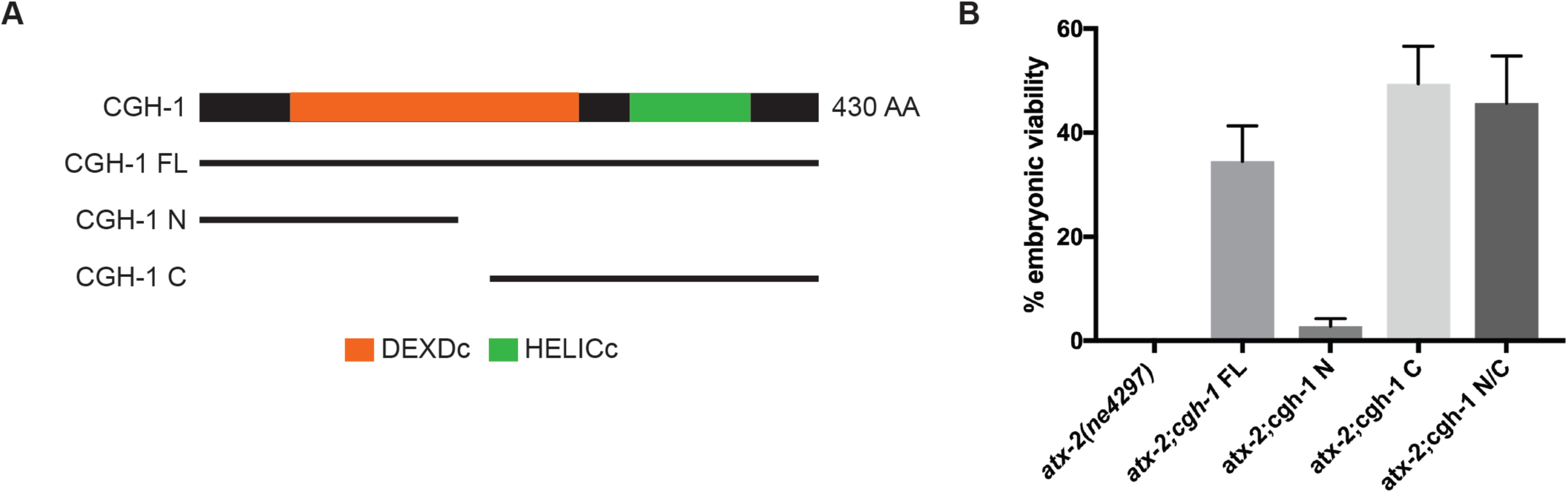
Alternative CGH-1 feeding RNAi targets still suppress *atx-2(ne4297)* (A-B) Three different feeding RNAi plasmids specific to different regions of CGH-1 all suppress the cell embryonic lethality and cell division defects observed in the *atx-2(ne4297)* suppression screen by the full length CGH-1 RNAi.

## Supplemental movies

**Movie S1.** Relating to Figure 2C. Control embryo undergoing first cell division.

**Movie S2.** Relating to Figure 2C. Cytokinesis failure in an *cgh-1* fRNAi-treated embryo

**Movie S3.** Relating to Figure 2C. Cytokinesis fails in an *atx-2 (ne4297)* embryo at the restrictive temperature (24°C).

**Movie S4.** Relating to Figure 2C. An example of cytokinesis failure in a *cgh-1* fRNAi-treated *atx-2 (ne4297)* embryo at the restrictive temperature (24°C).

**Movie S5.** Relating to Figure 2C. An example of cytokinesis rescue in a *cgh-1* fRNAi-treated *atx-2 (ne4297)* embryo at the restrictive temperature (24°C).

**Movie S6.** Relating to Figure 3A. Control embryo expressing GFP-PAR-5 undergoing first cell division.

**Movie S7.** Relating to Figure 3A. *atx-2* fRNAi-treated embryo expressing GFP-PAR-5 undergoing first cell division.

**Movie S8.** Relating to Figure 3A. *cgh-1* fRNAi-treated embryo expressing GFP-PAR-5 undergoing first cell division.

**Movie S9.** Relating to Figure 3A. *atx-2; cgh-1* fRNAi-treated embryo expressing GFP-PAR-5 undergoing first cell division.

